# Deep Learning Models for Atypical Serotoninergic Cells Recognition

**DOI:** 10.1101/2024.03.03.583157

**Authors:** Daniele Corradetti, Alessandro Bernardi, Renato Corradetti

## Abstract

**Background:** The serotonergic system modulates brain processes via functionally distinct subpopulations of neurons with heterogeneous properties, including their electrophysiological activity. In extracellular recordings, serotonergic neurons to be investigated for their functional properties are commonly identified on the basis of “typical” features of their activity, i.e. slow regular firing and relatively long duration of spikes. Thus, due to the lack of equally robust criteria for discriminating serotonergic neurons with “atypical” features from non-serotonergic cells, the physiological relevance of the diversity of serotonergic neuron activities results largely understudied.

**New Methods:** We propose deep learning models capable of discriminating typical and atypical serotonergic neurons from non-serotonergic cells with high accuracy. The research utilized electrophysiological in vitro recordings from serotonergic neurons identified by the expression of fluorescent proteins specific to the serotonergic system and non-serotonergic cells. These recordings formed the basis of the training, validation, and testing data for the deep learning models. The study employed convolutional neural networks (CNNs), known for their efficiency in pattern recognition, to classify neurons based on the specific characteristics of their action potentials.

**Results:** The models were trained on a dataset comprising 43,327 original spike samples, alongside an extensive set of 6.7 million synthetic spike samples, designed to mitigate the risk of overfitting the background noise in the recordings, a potential source of bias. Results show that the models achieved high accuracy and were further validated on “non-homogeneous” data, i.e., data not used for constructing the model, to confirm their robustness and reliability in real-world experimental conditions.

**Comparison with existing methods:** Conventional methods for identifying serotonergic neurons allow recognition of serotonergic neurons defined as typical. Our model based on the analysis of the sole spike reliably recognizes over 92 features of spike and activity.

**Conclusions:** The model is ready for use in experiments conducted with the here described recording parameters. We release the codes and procedures allowing to adapt the model to different acquisition parameters or for identification of other classes of spontaneously active neurons.

**PACS:** 87.19.L, 87.19.lv, 87.85.dm, 07.05.Mh, 87.85.Tu

## 1. Introduction

Activity of serotonergic neurons is known to regulate a wealth of autonomic and higher functions in mammals (Steinbusch et al., 2021; Faulkner and Deakin, 2014; Pilowsky, 2014; Lesch et al., 2012; Monti, 2011). Present knowledge of the physiological and pharmacological properties of serotonergic neurons is mostly based on electrophysiological recordings of neuronal activity from raphe nuclei of laboratory animals both *in vivo* and *in vitro*. However, most of recordings have been performed on neurons whose serotonergic identity was based on criteria that were empirically developed in the years to restrict the investigations to recordings from neurons that displayed very typical activity. For serotonergic neurons, the accepted criteria require the concomitant regularity of firing, broad action potential and, when pharmacological assays were allowed by the experimental design, sensitivity to serotonin1A receptor agonists that typically produce reversible slowing or cessation of neuron firing. When recordings are conducted in slices under microscopy guidance, the large size of serotonergic neuron soma could be used as an additional criterion. Adhering to these strict criteria for serotonergic neuron identification, however, results in a selection bias that has limited the studies to the “typical” neurons which might underrepresent the variety of serotonergic neuron population. Indeed, evidence for the existence of subpopulations of serotonergic neurons with distinctive neurochemical and pharmacological properties as well as firing patterns, emerged in the course of the past 40 years of dedicated research (e.g. Calizo et al., 2011; Paquelet et al., 2022; see also in Gaspar et al., 2012; Andrade and Haj-Dahmane, 2013; Commons, 2020). For instance, using *in vitro* recordings from dorsal raphe nucleus the possibility that serotonergic neurons display also irregular firing or peculiar rhythmic fluctuations in firing activity has been described since early recordings both *in vivo* and *in vitro* (Mosko and Jacobs (1974, 1976) and more recently confirmed with recordings of serotonergic neurons from transgenic mice selectively expressing fluorescent proteins in serotonergic neurons (Mlinar et al, 2016). Thus, the principal drawback of the intra-experiment recognition of 5-HT neurons is that serotonergic neurons displaying atypical activity or spikes narrower than expected are discarded and their pharmacological and physiological characteristics remain elusive. In addition, in the course of our research on genetically fluorescent serotonergic neurons (Montalbano et al., 2015; Mlinar 2016) we also noticed the existence of non-serotonergic (non-fluorescence labelled) neurons with regular activity and relatively broad spikes whose duration often overlaps that of spikes recorded in serotonergic neurons. Thus, in “real life” experimental conditions the activity characteristics of a non-neglectable number of serotonergic and non-serotonergic neurons could overlap and adherence to the above-mentioned strict criteria for identification of typical serotonergic neurons has the advantage to ensure a reasonable homogeneity of the population under study, in spite of the selection bias introduced. On the other hand, the characteristics of what we define “atypical” serotonergic neurons remain understudied.

In the present work we have taken advantage of the recordings present in our internal database and obtained from transgenic mice selectively expressing fluorescent proteins in serotonergic neurons to develop deep-learning based models for recognition of serotonergic and non-serotonergic neurons with relatively high accuracy and that can be implemented in the recording programs to quickly help the experimenter in the decision of continuing the recording or to change the experimental design, should an atypical serotonergic or non-serotonergic neuron be identified.

## 2. Material and Methods

### 2.1. Source database

To train, test and validate our deep-learning based models we used the original recordings from our internal database built in the occasion of our studies in which we described the firing characteristics of genetically identified dorsal raphe serotonergic neurons in brain slices. Serotonergic and non-serotonergic neurons were thus identified on the basis of a parameter independent from their electrophysiological features, i.e.on serotonergic system-specific fluorescent protein expression (serotonergic) or lack of expression (non-serotonergic). In our original articles (Mlinar et al., 2016; Montalbano et al., 2015) we detailed the procedure to obtain the three transgenic mouse lines with serotonergic system-specific fluorescent protein expression used in the present work: Tph2::SCFP; Pet1-Cre::Rosa26.YFP ; Pet1-Cre::CAG.eGFP.

### 2.2. Loose-seal cell-attached recordings

Detailed description of the electrophysiological methods and of the measures for improving reliability of loose-seal cell-attached recordings has been previously published (Montalbano et al., 2015; Mlinar et al., 2016). In brief, mice (4-28 weeks of age) were anesthetized with isofluorane and decapitated. The brains were rapidly removed and dissected in ice-cold gassed (95% O2 and 5% CO2) ACSF composed of: 124 mM NaCl, 2.75 mM KCl, 1.25 mM NaH2PO4, 1.3 mM MgCl2, 2 mM CaCl2, 26 mM NaHCO3, 11 mM D-glucose. The brainstem was sliced coronally into 200 µm thick slices with a vibratome (DSK, T1000, Dosaka, Japan). Slices were allowed to recover for at least 1 h at room temperature and then were individually transferred to a submersion type recording chamber and continuously superfused at a flow rate of 2 ml min-1 with oxygenated ACSF warmed to 37°C by a feedback-controlled in-line heater (TC-324B / SF-28, Warner Instruments, Hamden, CT). Slices were allowed to equilibrate for 10-20 min before the beginning of the recording. To reproduce in brain slices noradrenergic drive that facilitates serotonergic neuron firing during wakefulness (Baraban and Aghajanian, 1980; Levine and Jacobs, 1992), ACSF was supplemented with the natural agonist noradrenaline (30 *µ*M) or with the *α*1 adrenergic receptor agonist phenylephrine (10 *µ*M; Vandermaelen and Aghajanian, 1983). Neurons within DRN were visualized by infrared Dodt gradient contrast video microscopy, using a 40X water-immersion objective (N-Achroplan, numerical aperture 0.75, Zeiss, Göttingen, Germany) and a digital CCD camera (ORCA-ER C4742-80-12AG; Hamamatsu, Hamamatsu City, Japan) mounted on an upright microscope (Axio Examiner Z1; Zeiss) controlled by Axiovision software (Zeiss). Loose-seal cell-attached recordings were made from fluorescent protein-expressing or not expressing neurons, visually identified by using Zeiss FilterSet 46 (eGFP and YFP, excitation BP 500/20, emission BP 535/30) or Zeiss FilterSet 47 (CFP, excitation BP 436/20, emission BP 480/40). Fluorescence was excited using a Zeiss HXP 120 lamp. Patch electrodes (3-6 MΩ) were pulled from thick-walled borosilicate capillaries (1.50 mm outer diameter, 0.86 mm inner diameter; Corning) on a P-97 Brown-Flaming puller (Sutter Instruments, Novato, CA) and filled with solution containing (in mM): 125 NaCl, 10 HEPES, 2.75 KCl, 2 CaCl2 and 1.3 MgCl2, pH 7.4 with NaOH. After positioning the pipette, development of loose-seal was monitored by using a voltage-clamp protocol with holding potential of 0 mV and test pulse of 1 mV / 100 ms, repeated every second. Weak positive pressure was released and gentle suction was slowly applied until detected spikes increased to 50 - 100 pA peak-to-peak amplitude. In some experiments this procedure was repeated during recording to increase signal to noise ratio.. Corresponding seal resistance was in 10 to 20 MΩ range. Recordings were made using an Axopatch 200B amplifier (Molecular Devices, Sunnyvale, CA) controlled by Clampex 9.2 software (Molecular Devices). Signals were low-pass filtered with a cut-off frequency of 5 kHz (Bessel) and digitized with sampling rate of 40 kHz (Digidata 1322A, Molecular Devices). After the recording, images of recorded neuron were acquired to document the expression of the fluorescent marker in the recorded neuron.

### 2.3. Offline Analysis of recordings

Detection of spikes was performed using event detection routine of Clampfit 9.2 software. Spike duration (width) was determined from the shape of averaged spike by measuring the interval between the spike upstroke and the downstroke (or second downstroke, whenever present) hereby named UDI (Upstroke-Downstroke Interval) for convenience (see Fig. 6; see also Fig. 3 in Mlinar et al., 2016).

## 3. A Deep Learning Model

Recognizing serotonergic cells is a binary classification problem, i.e., serotonergic vs. non-serotonergic cells, for which deep learning (DL) algorithms and, more specifically, the use of convolutional neural networks (CNN) have yielded excellent results. Notably, CNN are inspired by the organization of the animal visual system, particularly the human brain, and excel at tasks like image feature extraction, which is fundamental for recognition purposes (Liu, 2018). They employ mechanisms such as feedforward inhibition to alleviate issues like gradient vanishing, enhancing their effectiveness in complex pattern recognition tasks (Liu et al, 2019). With these considerations in mind, we have chosen to use a CNN architecture even in the apparently unconventional context of numerical pattern recognition, i.e., the recorded signal of a neuronal cell. The inspiring idea behind this choice is to leverage the ability of CNNs to amplify numerical patterns that occur at different scales, in this case within time intervals that are orders of magnitude smaller than the entire examined signal. In fact, this is a characteristic typical of neuronal spikes, where the maximum peak impulse can occur within a scale of 1 ms, while the firing period, i.e., the time interval between two consecutive spikes, can be two orders of magnitude greater.

### 3.1. Preliminary approaches and definition of appropriate parameters for developing the model

Starting from the assumption that two factors are typically relevant in recognizing serotonergic cells, namely the specific shape of the spike together with its repetitiveness and firing frequency, we initially decided to consider time segments of 7 seconds as training data for the neural network. This ensured an adequate number of spikes to evaluate their consistency and periodicity. After several attempts in this direction, however, we realized that the importance of the cell’s spike shape was so predominant that the information obtained from analyzing the firing periodicity alone was not sufficient to compensate for the accuracy gained by focusing on the individual spike.

Our first preliminary analysis was done on 108 serotonergic cells and 45 non-serotonergic cells. Every spike for the training consisted in the recording of 7 ms taken from 2 ms before the detection threshold to 5 ms after. While the final accuracy of the resulting models was fairly high, ranging from 94.3% to 99.3%, further analysis on non-homogenous data, i.e. data collected on experimental days different from those used for the training and evaluation of the models, showed a much lower accuracy, which was a strong sign of the overfitting. Further investigation allowed to identify the primary source of overfitting in the background noise of the recordings which, having a specific signature, the model learned to incorporate in the recognition of the neuron types. Thus, models trained with spikes embedded in 7 ms time-segments learned how to classify the spikes on the basis of the background noise instead of the peculiar shape of the event.

Therefore, we decided to reduce the impact of the background noise present in the samples by limiting the time-window of spike analysis to 4 ms. This solution worked well, since we had a comparable accuracy of the metrics on non-homogenous data.

To obtain an additional estimate of the impact of the background noise on models trained on 4 ms samples we produced an high volume of synthetic spike samples that would incorporate the original data mixed with different noise backgrounds. To this purpose, 43,327 original spikes we produced about 6.7M synthetic spikes (see below for a description of the generation process). In this new set of synthetic data, every spike had 150 different noise backgrounds, thus reducing the impact that such noise could have in the training. The performance of the additional model trained with these data was used to confirm that the impact of the background noise in 4 ms spike samples was negligible.

### 3.2. Data used for originating and validating the final model

*Original Training Data: .* The original data for the training, validation and testing of the models consisted in 43,327 spike samples extracted from 108 serotonergic cells and 45 non-serotonergic cells. More specifically, we extracted 29,773 spikes from serotonergic cells, and 13,554 spikes from nonserotonergic cells. In all cases, the triggering threshold of the event was -50 pA and the spike was then sampled 1 ms before the triggering threshold until 3 ms after (see Fig. 1). Since the sampling rate of the original recordings was 40 kHz, every spike sample consists of 160 values. All the samples were then randomly subdivided into 30,328 for training, 6,500 for validation and 6,499 for testing.

**Figure 1:**
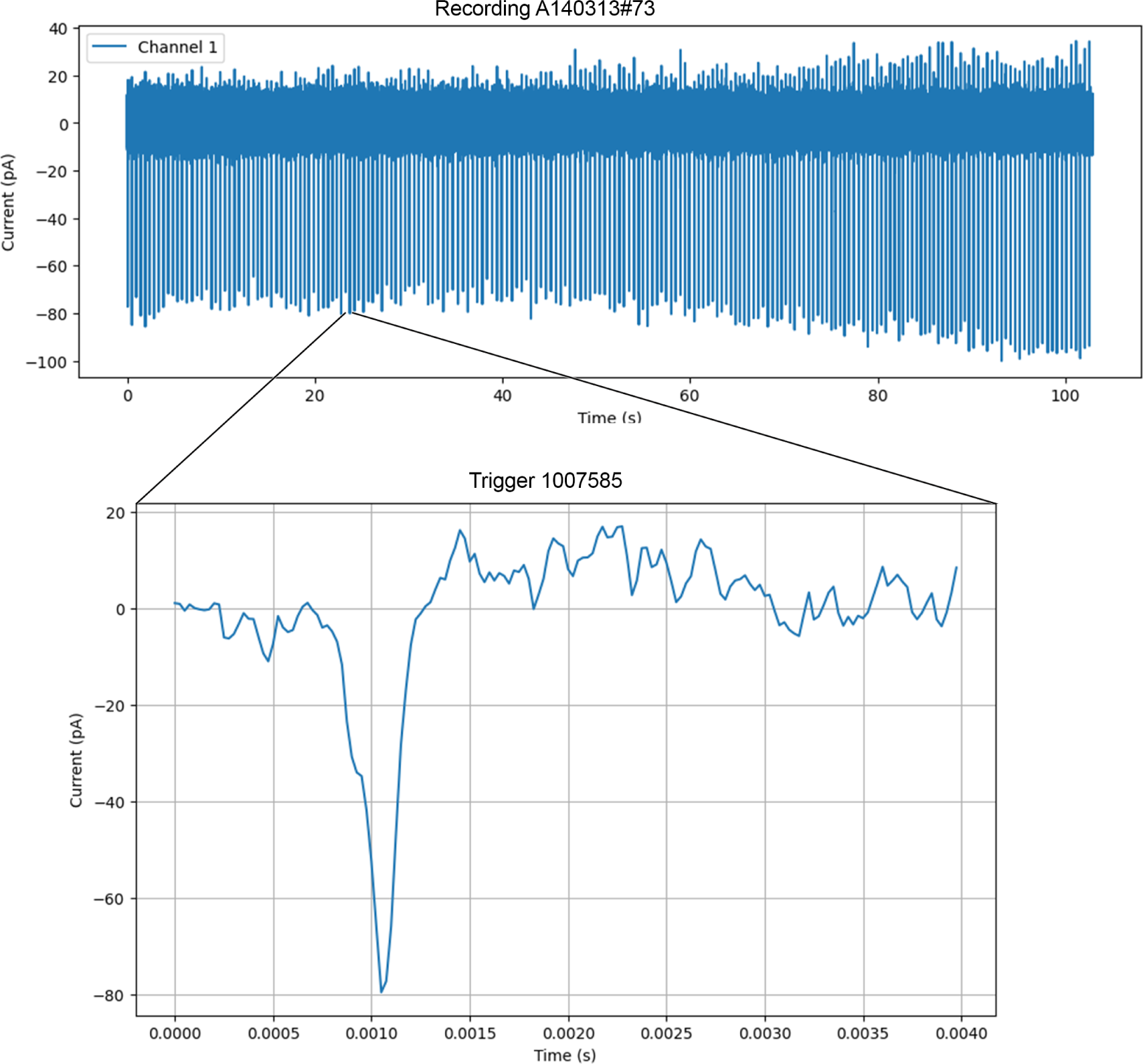
Example on how single events were isolated and selected. The image depicts the recording of the serotonergic cell A140313#073 and the 4 ms event of triggered at point 1007585, i.e. at 25.189 sec.

*Non-homogenous Data:.* The non-homogenous data consisted in 24,616 samples extracted from 55 serotonergic cells (18,595 spikes) and 26 non-serotonergic cells (6,021 spikes) collected in experimental days not used to obtain the training data, thus with different signal noise. These data were never part of the training set, nor validation, nor testing set during the training, and constituted just an additional independent test for the already trained model.

*Synthetic Data:.* The synthetic data consisted in 6,675,300 spike samples of 160 points (simulating 4 ms at 40 kHz of sampling) arising from the 43,327 original training data samples. From the original training data recordings we extracted 600 noise masks (see e. g. Fig. 2) from which 150 were randomly applied to the 43,327 spikes thus obtaining the synthetic data (see e. g. Fig. 3).

**Figure 2:**
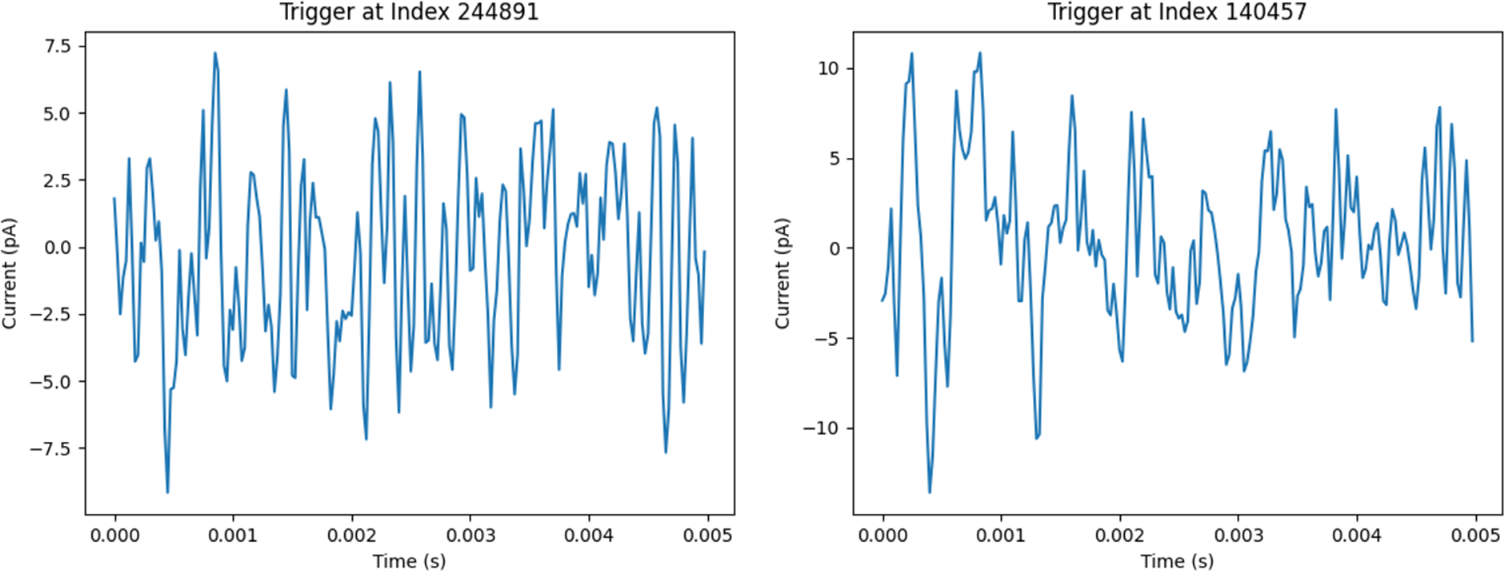
Examples of noise masks collected from the recordings of cell A140724#065 (*on the left*) and A160127#015 (*on the right*).

The generation of the synthetic data was done according the following procedure. Each original training data sample is smoothed through averaging, i.e. the values of the smoothed sample 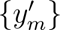 with *m* ∈ {1, …, 160}, are given as the averages of the values of the original sample {*y_m_*} by

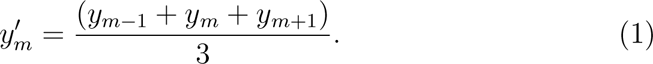

Then the values of smoothed spike 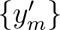 are added to the values of randomly chosen noise mask 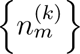 where *k* ∈ {1*, …,* 600} is randomly chosen. The final synthetic sample is thus obtained as the sample {*y*^(^*^k^*^)^*_m_*} with

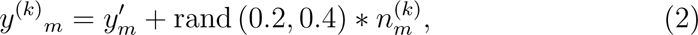

where rand (0.2, 0.4) is a randomly generated coefficient around 1/3 to modulate the noise.

### 3.3. Model Description

In accordance with the origin of our dataset, we developed two distinct models, namely the “biological” model (trained only over original data) and the “synthetic” model (trained only on synthetic data). The biological model underwent training, validation, and testing using the original training data, which comprises 43,327 spike samples. Conversely, the synthetic model was trained, validated, and tested utilizing synthetic data, encompassing 6,675,300 spike samples. We emphasize that, in the present experimental context, the use of the synthetic model was exclusively for monitoring the overfitting phenomena and noise biases in the classification process of the biological model.

Fig. 4 summarizes the various steps used to implement the model from recorded signals. The architecture of the models is a sequence of layers commonly used in deep learning, specifically in the context of convolutional neural networks (CNNs) for image or signal processing. We implemented the architecture using the Keras libraries in TensorFlow 2. The model of the neural network consists of a normalization layer for stabilizing the learning process and reducing training time; two repetitions of a 2D convolutional layer with 32 filters and a max pooling layer with a pool size of (2x1); a flatten layer to connect to a dropout layer and dense layers with 2 output units used for binary classification. Activation functions of the convolutional layers are the ReLU, while for the dense layer we used the classic sigmoid (see Table 1 for a summary of the model). For training we chose the “binary crossentropy” loss function, which is standard for binary classification problems, while the optimizer was “Adam” (Adaptive Moment Estimation) as these are common choices. A special treatment was devoted to the kernel of the 2D convolutional layers. Indeed, since the kernel of these layers express the ability of the convolutional process in enlarging a specific portion of the pattern, we explored a range of possible kernels between 1 to 32. Apart from the models with kernel *<*5, the results were homogenous and summarized in Fig. 5. All models were trained on 25 epochs with a batch size of 64 and their test accuracy ranged from 90.6% (model with kernel 1) to 98.7% (model with kernel 23, 24 and 27) with a test loss of 0.2221 and 0.037106. To enhance the robustness of the model, instead of selecting a single kernel and using one model for inference, we selected all models with kernels ranging between 20 and 30 and took the consensus between the models. This technique ensures more stability in the overall architecture and is often considered best practice.

**Figure 3:**
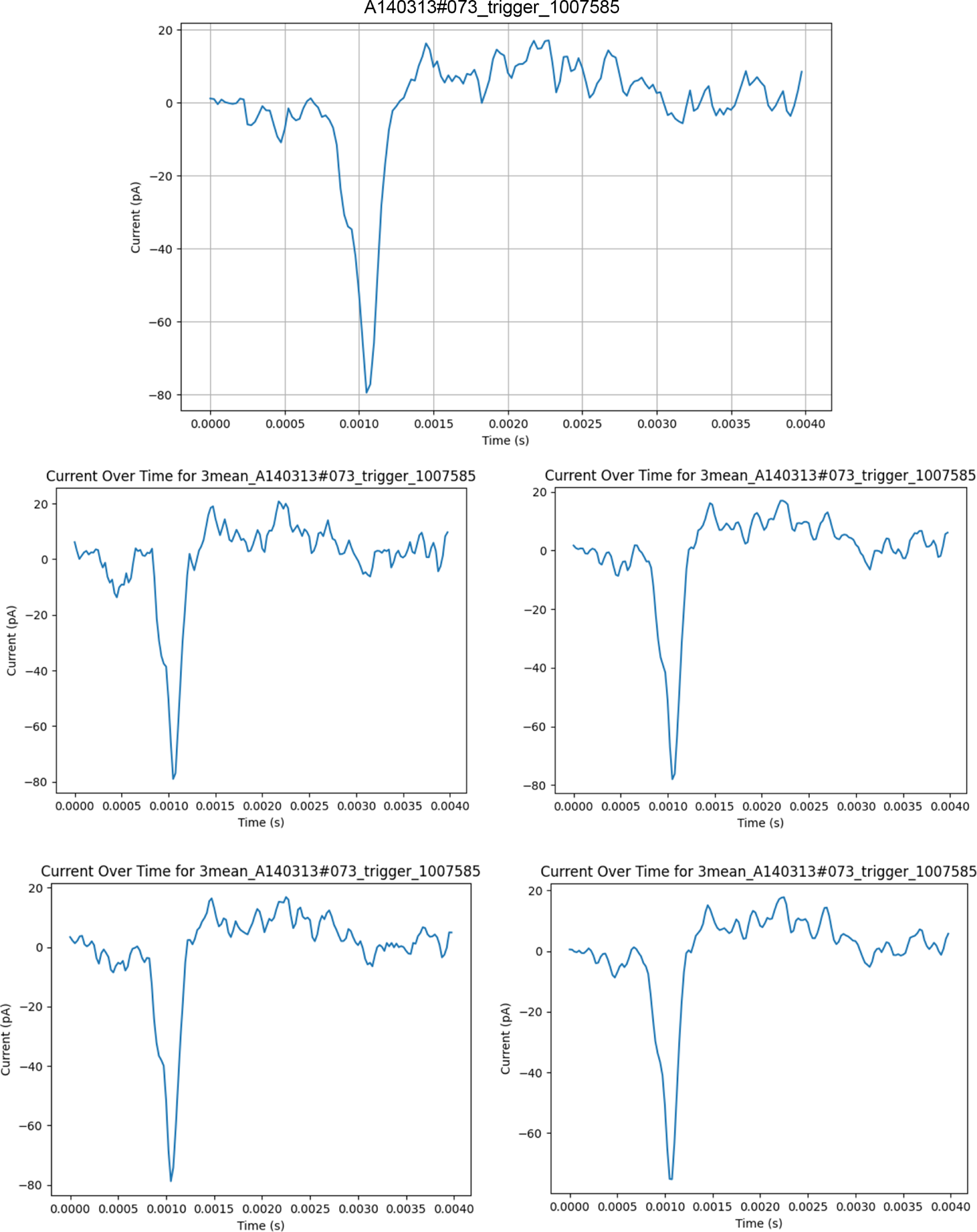
Example of 4 synthetic spikes generated by the event triggered at 1007585, i.e. 25.189 sec, of the serotonergic cell A140313#073. Top trace: the original recording of the event. The panels report four spike obtained by processing the original trace with different noise masks (see methods).

**Figure 4:**
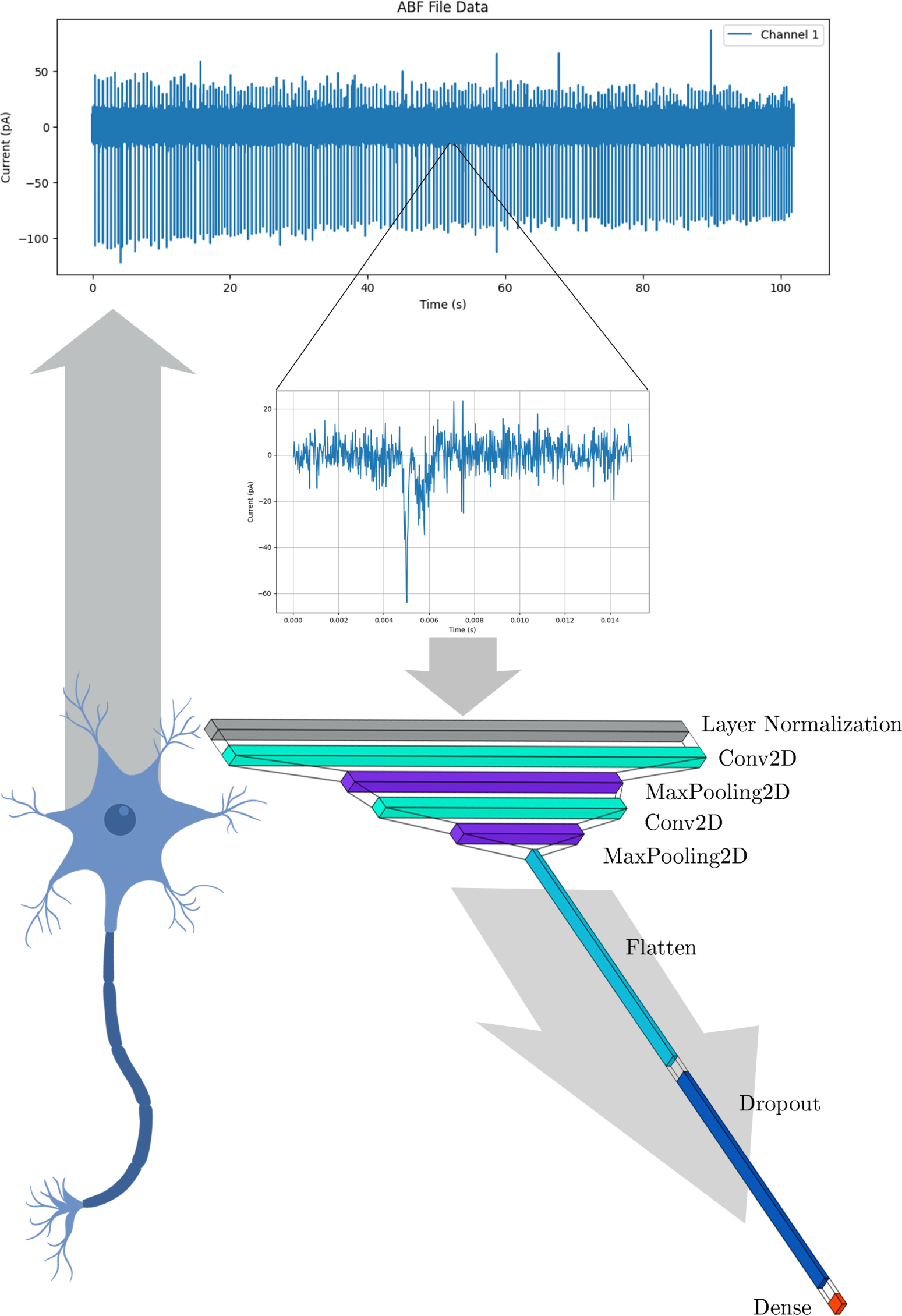
Summary of the various steps used to implement the model from recorded signals: from neuronal cell the signal is sam1p1led at 40 kHz and recorded as .abf file, then the all events are selected and sent to 10 neural networks with the above architecture for classification (only difference between the architectures is the value of the 2D convolutional kernel with ranges between 20 to 30).

**Figure 5:**
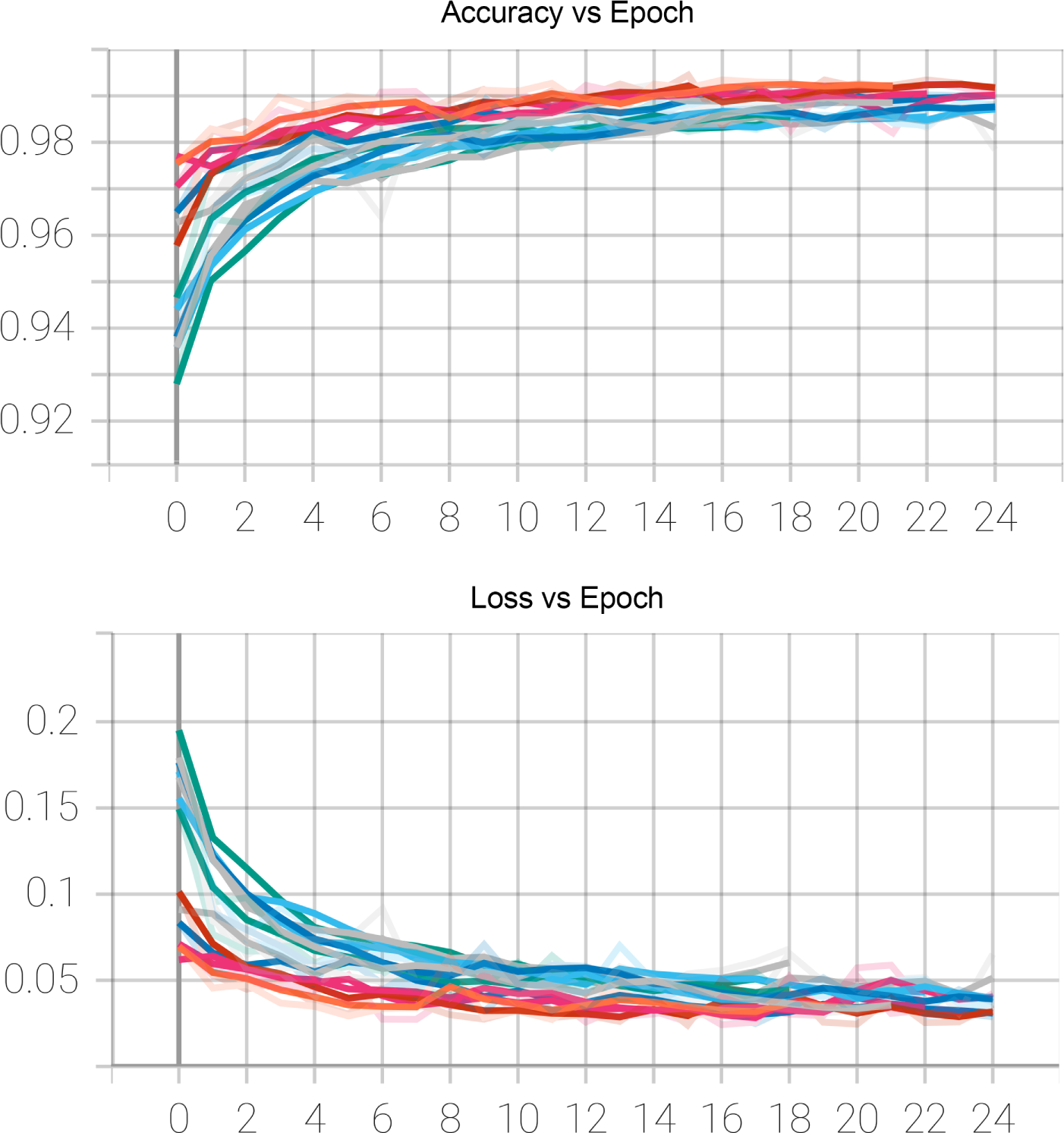
*On top*: in the graphic is depicted the accuracy for every epoch of the training (0-24) for all models with kernel ranging from 20 to 30. *On the bottom*: the image depicts the loss function for every epoch of the training (0-24) for all models with kernel ranging from 20 to 30.

**Table 1:** Summary of the CNN architectural model with kernel 20 used for the neural network. Other models follow the same architectural structure and change only for the dimension of the kernel.

| *Layer (type)* | *Output Shape* | *Param #* |
| --- | --- | --- |
| Layer Normalization | (None, 160, 2, 1) | 320 |
| Conv2D | (None, 141, 2, 32) | 672 |
| MaxPooling2D | (None, 70, 2, 32) | 0 |
| Conv2D | (None, 51, 2, 64) | 41024 |
| MaxPooling2D | (None, 25, 1, 64) | 0 |
| Flatten | (None, 1600) | 0 |
| Dropout | (None, 1600) | 0 |
| Dense | (None, 2) | 3202 |
| *Total Params* | 45218 |  |

Finally, it is worth noticing that while the training of the biological model did not required any specific adjustment, the synthetic model, involving *>* 6M spike samples required a continuous learning implementation, where the model was trained over 105 training sessions of 63,450 synthetic spike samples.

### 3.4. Assessment of Accuracy and Sensitivity

For the assessment of the models we used the following metrics: *Accuracy*, *Sensitivity at Specificity 0.5* and the *Confusion Matrix*.

- Accuracy measures the proportion of total predictions (both serotonergic and non-serotonergic cells) that the model correctly identifies, i.e.

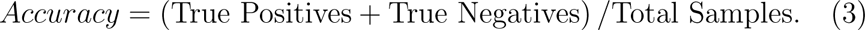

This metric was chosen for identifying if the models are generally effective in classifying both serotonergic and non-serotonergic cells.

- Sensitivity at Specificity measures the sensitivity of the model, i.e.

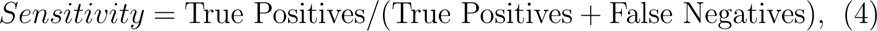

at a fixed specificity, i.e. True Negatives/(True Negatives + False Positives), which we set at 0.5. The choice of this metric with this setting ensures that the models are not overly biased towards identifying serotonergic cells at the expense of misclassifying non-serotonergic ones.

- Finally, the Confusion Matrix shows the percentages of True Positives, False Positives, True Negatives, and False Negatives giving a complete feedback of the models. This is a a detailed view that we considered essential for understanding the specific areas where the models need improvements.

All these metrics were used for all the data, i.e. Original Training Data, Nonhomogenous Data and Synthetic Data. In the specific case of the Original Training Data all the metrics were used in the three phases of Training, Validation and Testing. The training phase was developed on 30,328 spikes selected uniquely for training. The Validation phase, which is used to tune hyperparameters, was on 6,500 spikes which the model has not seen during training. Finally, the last 6499 spikes were used for the Testing of the models, and are those on which the true performance of the models is assessed.

### 3.5. Repository of the Model and Data

We made available in the github respository (Corradetti et al., 2024) the following:

1. the .abf recordings of original training data and the non-homogenous data,
2. the 43,327 single spikes samples of the orginal training data stored in .csv files of 160 points,
3. the 24,616 single spikes samples of the non-homogenous data stored in .csv files of 160 points,
4. the 6,675,300 million single spikes samples of the synthetic data stored as numpy vector,
5. the trained models with different kernels,
6. the results of the models,
7. the Python notebooks for training of the models and for inference.

## 4. Results

In this study we compared the spiking activity of 300 neurons recorded in DRN slices obtained from transgenic mouse lines with serotonergic system-specific fluorescent protein expression.

### 4.1. Visual discrimination of spikes

As illustrated in Fig. 6, serotonergic neurons displayed spikes of different shape and duration that were often difficult to be discriminated from those observed in non-serotonergic neurons. Thus, with the exception of spikes showing the typical shape and duration of serotonergic neurons (e.g. Fig. 6: traces a1, a2) or of non-serotonergic cells (e.g. Fig. 6: trace b1) both types of neurons may display spikes similar in width and/or shape. Therefore, the sole duration of the spike, which could be determined online by measuring the upstroke/downstroke interval (UDI) may result not enough indicative for immediate serotonergic neuron identification.

**Figure 6:**
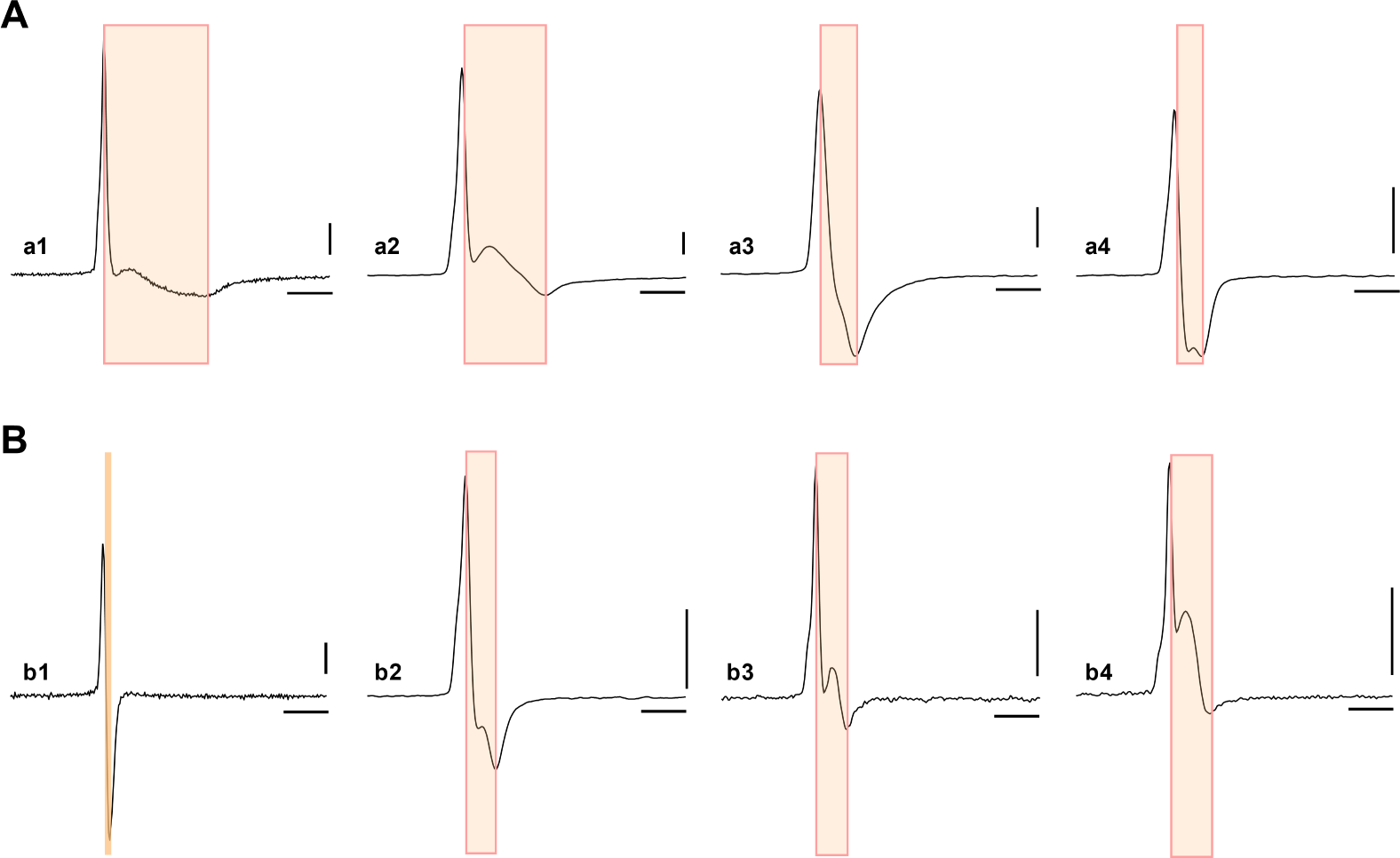
Examples of spikes recorded from serotonergic and non-serotonergic neurons in slices of dorsal raphe nucleus. A. Fluorescent protein-labelled (serotonergic) neurons: a1,a2: typical spikes of serotonergic neurons; note the long interval between spike upstroke and downstroke (UDI) highlighted by the shaded area in all traces. a3-a4: recordings from serotonergic neurons displaying spikes of shorter duration. B. Fluorescent proteinunlabelled (non-serotonergic) neurons: b1: typical biphasic spike of short duration from a non-serotonergic neuron; b2-b4: spikes of variable shape recorded from non-serotonergic neurons. Shaded areas indicate the width of the spike measured by UDI (see methods). Note the overlap in spike width of some serotonergic and non-serotonergic neurons. Traces are averages of 15-50 sweeps. Calibrations 25 pA (polarity inverted); 1 ms.

From our database of recordings we have selected 150 serotonergic neurons labelled by fuorescent proteins and 150 non labelled cells, deemed to be nonserotonergic cells. The distribution of spike width of these two populations is shown in Fig. 7. These neurons were chosen on the sole technical characteristic of not showing detectable artefactual transients that could be mistaken by the deep-learning routine as action potentials. From these two populations of neurons we extracted 108 serotonergic and 45 non-serotonergic neurons to implement the training of the Biological Model. In addition, 12 serotonergic neurons from three different experimental days and 10 non-serotonergic neurons from four different experimental days were used for testing the model with data non homogeneous to the training (see methods).

**Figure 7:**
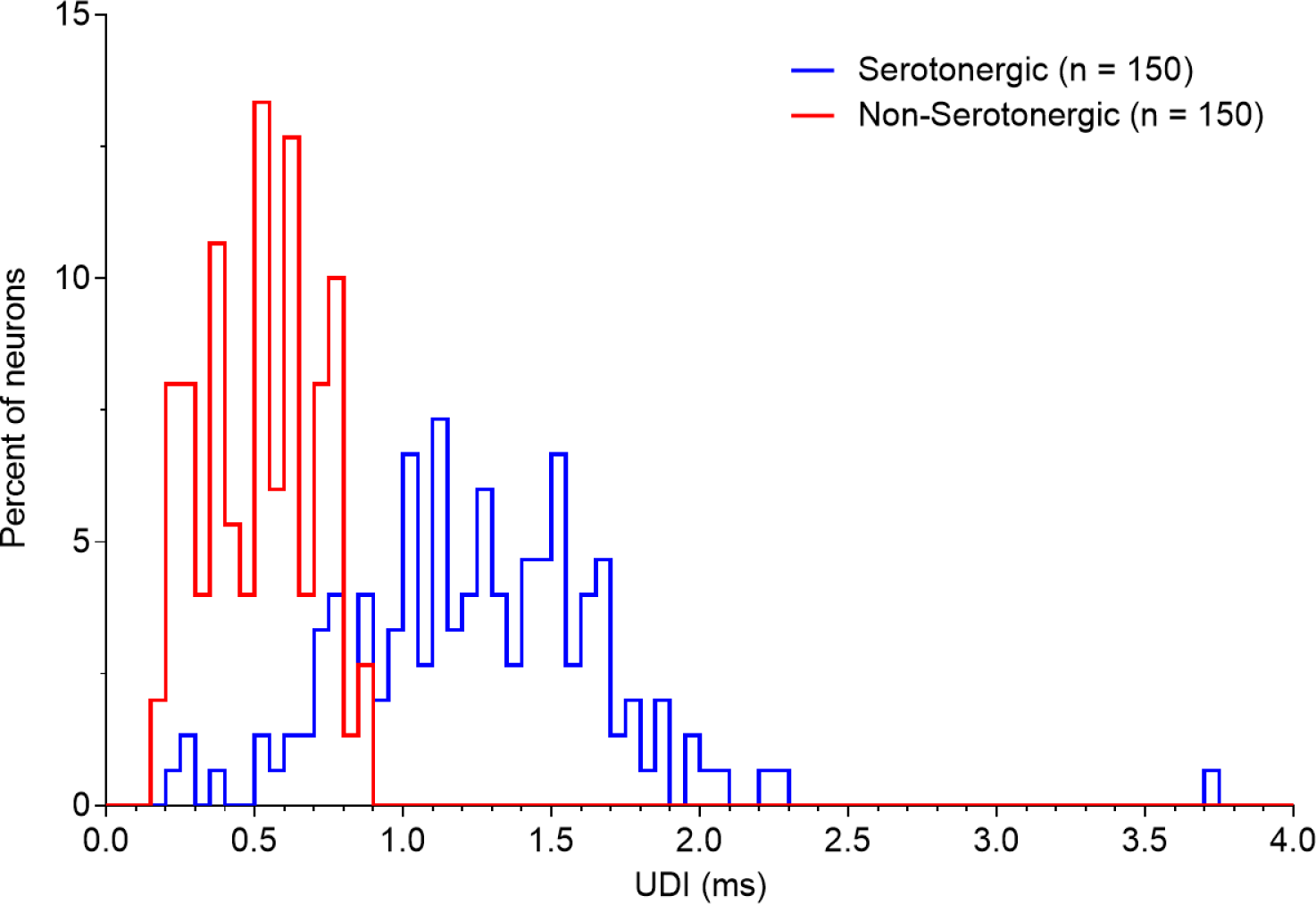
Distribution of spike duration of serotonergic and non-serotonergic neurons. Histograms report the distribution of spike width measured by the interval between spike upstroke and downstroke (UDI) in serotonergic (blue) and non-serotonergic neurons (red) recorded in slices of dorsal raphe nucleus. Note the overlap in spike duration between serotonergic and non-serotonergic neurons.

As shown in Fig. 8, the neurons used from training and testing the model 1 are representative of the two (serotonergic and non-serotonergic) populations of neurons.

**Figure 8:**
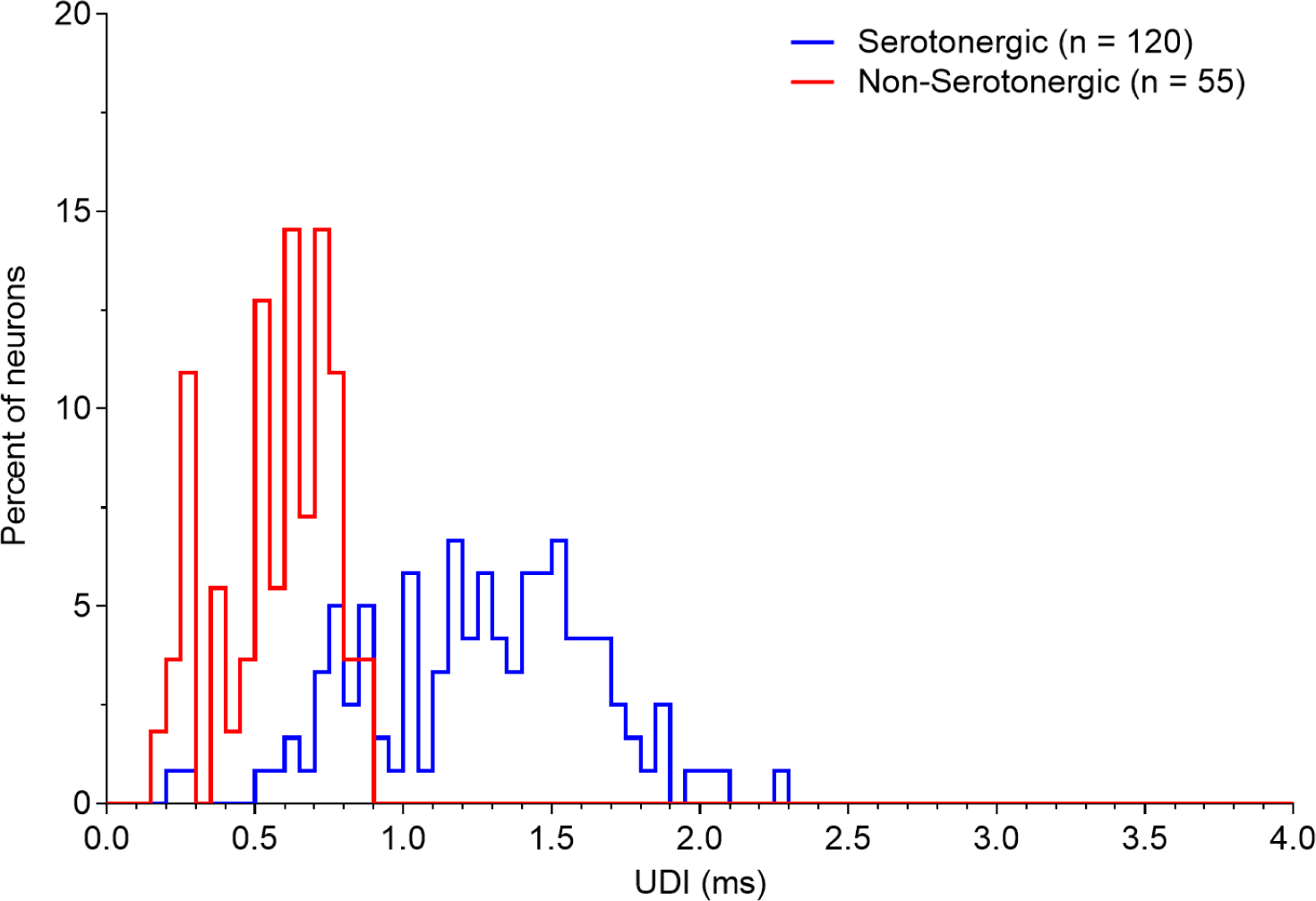
Distribution of spike duration recorded from the neurons utilized to develop the model 1. Histograms report the distribution of spike width measured by the interval between spike upstroke and downstroke (UDI) in serotonergic (blue) and non-serotonergic neurons (red) recorded in slices of dorsal raphe nucleus.

An additional group of recordings (*n* = 30) from fluorescence identified serotonergic and non-serotonergic neurons, not previously used for the model implementation, were processed by the model 1 to test its ability to recognize cell type from the spike characteristics distilled by the model itself.

### 4.2. Discrimination with Deep Learning Models

The metrics of both the biological model and the synthetic model were collected over the testing data (original and synthetic) during their training phase, as standard practice in deep learning. Over this data both the biological model and the synthetic model scored *>* 98.5% accuracy. In addition to the standard practice we evaluated the models over non-homogenous data in order to evaluate possible sources of overfitting arising from noise signatures in the recordings. On this dataset both the biological model scored *>* 92.5% accuracy showing the existence of some light source of overfitting. Nevertheless, the results from the synthetic data, similar in result to the original data, suggest that the source of overfitting is not depending on the noise signature of the record. Overall, we consider the metrics evaluated on non-homogenous data more indicative and reliable than those arising from the training data.

#### 4.2.1. Results on the Training Data

*Biological Model.* The biological models, when tested on the original training dataset, showed varying performance metrics. For kernels ranging from 1 to 28, the test loss was observed between 0.222171 (kernel 1) and 0.035998 (kernel 28). Accuracy measurements ranged from 0.906462 (kernel 1) to 0.987692 (kernel 27), and sensitivity at a specificity of 0.5 varied from 0.996154 (kernel 1) to 0.999846 (kernel 9), as detailed in Fig. 9. The consensus biological model, derived from kernels 20 to 30, recorded a test loss of 0.043076, an accuracy of 0.985161, and a sensitivity at specificity 0.5 of 0.999147, as shown in Table 2).

**Figure 9:**
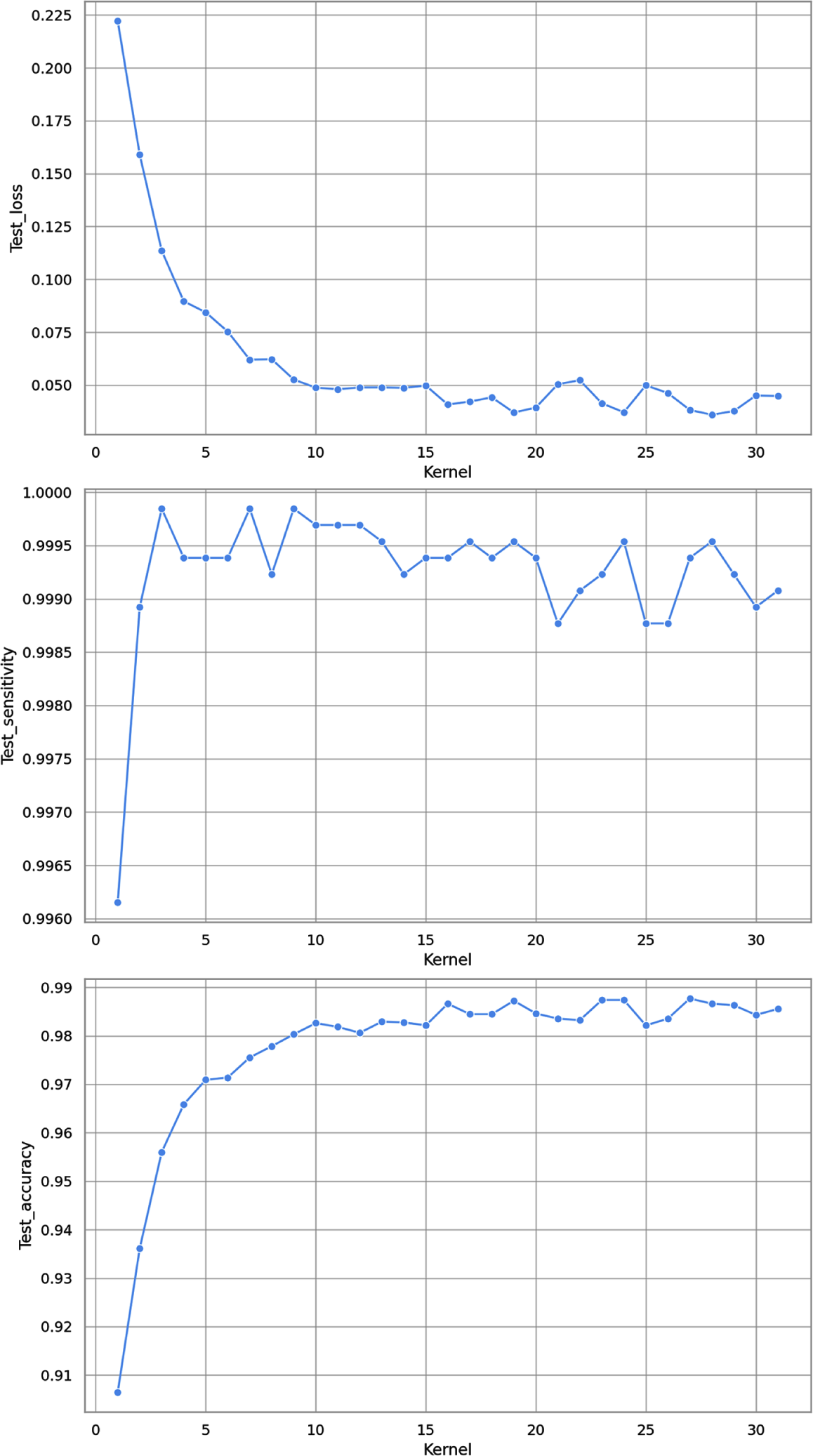
Each of the 32 models, with kernel sizes varying from 1 to 31, was evaluated for test loss, sensitivity at a specificity of 0.5, and accuracy. The resulting graphs depict a monotonic trend correlating with the increasing kernel sizes, which eventually stabilizes in the range from kernel size 20 to 31.

**Table 2:**
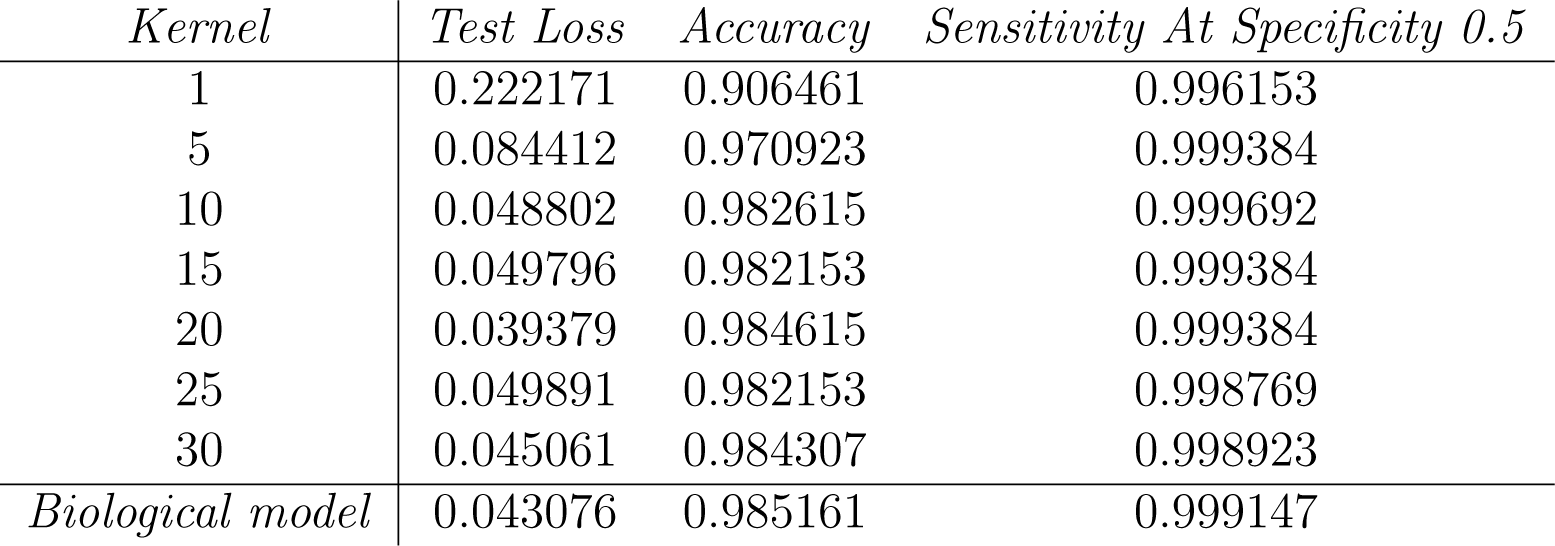
A selection of the metrics on the test data of the models trained with Original Training Data. Beside *Accuracy* and *Sensitivity At Specificity 0.5*, we reported also *Test loss*, which represent the error between the predicted values and the actual values and is a standard metric in evaluating DL models. Values reported in last row “*biological model*” refer to the metrics of the full biological model given as the consensus of the single models with 20 *≤* kernel *≤* 30.

| *Kernel* | *Test Loss* | *Accuracy* | *Sensitivity At Specificity 0.5* |
| --- | --- | --- | --- |
| 1 | 0.222171 | 0.906461 | 0.996153 |
| 5 | 0.084412 | 0.970923 | 0.999384 |
| 10 | 0.048802 | 0.982615 | 0.999692 |
| 15 | 0.049796 | 0.982153 | 0.999384 |
| 20 | 0.039379 | 0.984615 | 0.999384 |
| 25 | 0.049891 | 0.982153 | 0.998769 |
| 30 | 0.045061 | 0.984307 | 0.998923 |
| *Biological model* | 0.043076 | 0.985161 | 0.999147 |

*Synthetic Model.* The evaluation of the 32 synthetic models on the synthetic dataset yielded superior metrics compared to the biological models, as detailed in Table 3. However, these results are not deemed highly significant, as overfitting not related to recording noise tends to be amplified in the augmented dataset. Further analysis revealed that on non-homogeneous data, the performance of the synthetic models was on par with that of the biological models.

**Table 3:** A selection of the metrics on the test data of the models trained with synthetic data. Values reported in last row “*synthetic model*” refer to the metrics of the full synthetic model given as the consensus of the single models with 20 *≤* kernel *≤* 30. It is important to note that the synthetic model’s primary purpose is to monitor overfitting arising from the specific noise signature in the recording. Due to its heightened sensitivity to various other forms of overfitting, the metrics for this model, though provided for completness, are not considered relevant.

| *Kernel* | *Test loss* | *Accuracy* | *Sensitivity At Specificity 0.5* |
| --- | --- | --- | --- |
| 1 | 0.001323 | 0.999580 | 1 |
| 5 | 0.002248 | 0.999475 | 0.999895 |
| 10 | 0.002121 | 0.999685 | 0.999895 |
| 15 | 0.002424 | 0.999685 | 0.999895 |
| 20 | 0.002946 | 0.998741 | 1 |
| 25 | 0.000378 | 0.999790 | 1 |
| 30 | 0.000667 | 0.999895 | 1 |
| *Synthetic model* | 0.001196 | 0.999590 | 0.999971 |

#### 4.2.2. Results on Non-Homogenous Data

The most significant outcomes were derived from non-homogeneous data, i.e., cells that were not utilized in training and were collected on different days than the training data. Using this dataset, the biological model achieved an accuracy of 0.925 and a sensitivity at specificity of 0.5 of 0.88461. A crucial indicator of performance is the biological model’s confusion matrix (refer to Fig. 10). Out of 55 serotonergic cells, 52 (94.4%) were accurately identified as serotonergic (True Positive), while 3 (5.6%) were incorrectly classified as non-serotonergic (False Negative). Conversely, of the 26 non-serotonergic cells, 23 (88.4%) were correctly recognized (True Negative), and 3 (11%) were erroneously labeled as serotonergic (False Positive).

**Figure 10:**
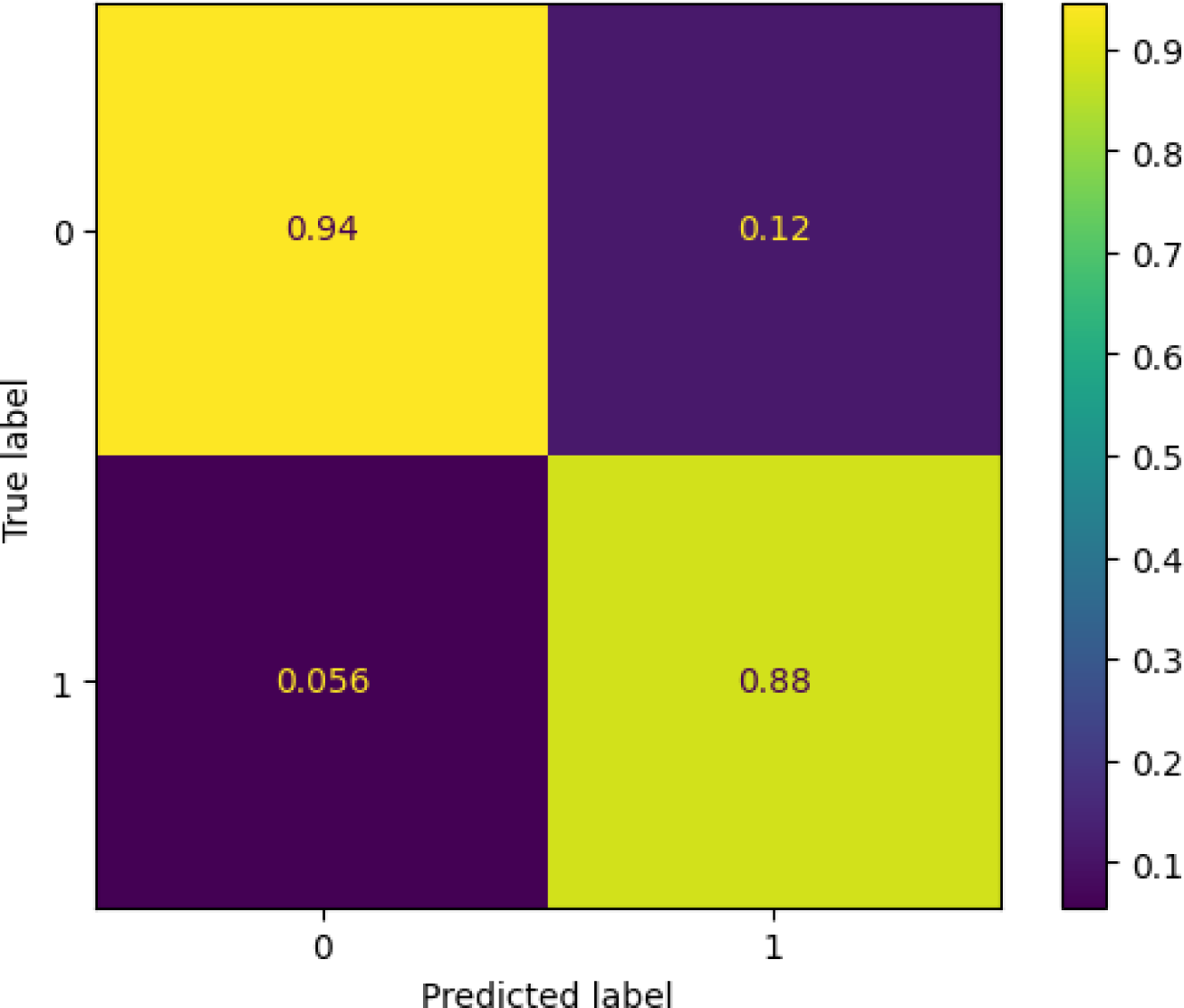
The confusion matrix for the biological model over the non-homogeneous data labels serotonergic cells as 0 and non-serotonergic cells as 1. The matrix shows the True Positive Rate 94.4% at the Top-Left; the False Negative Rate 5.6% at the Bottom-Left; the False Positive Rate 11% at the Top-Right; and the True Negative Rate 88.4% at the Bottom-Right.

Regarding the synthetic data, the confusion matrix values vary with each training session. The highest values attained by the synthetic data match those of the original data, specifically a 94.4% True Positive Rate, 5.6% False Negative Rate; 88.4% True Negative Rate and 11% False Positive Rate. The average values for the synthetic data across training sessions are 91.2% for True Positive Rate, 0.8% for False Negative Rate; 87.6% for True Negative Rate and 12.3% for False Positive Rate. These Fig.s for the synthetic data closely align with those of the original data, indicating no overfitting specifically due to the noise in the recorded signal.

## 5. Discussion

Deep-learning based models have gained increasing importance in biomedicine for their high performance in image processing and morphological recognition of cells that can be applied both in clinical diagnostics (Johansen et al., 2016; Litjens et al., 2017; Racz et al., 2020) and in preclinical research when complex patterns of data need to be measured, classified and interpreted (De Luca et al., 2023). More specifically, convolutional neural networks (CNN) effectively address complex pattern recognition especially when patterns are hidden across varying scales and orders of magnitude. This is highly relevant in neuronal spikes, where the peak impulse and the rise of the spike may occur in a fraction of millisecond, whereas the interval between spikes can be vastly longer. The here proposed model provides an important proof of concept for usefulness of CNN for identification of neuron types in the central nervous system on the basis of their spiking activity. To the best of our knowledge, this is the first time that this type of architecture is applied to recognition of neuronal spikes by their recorded traces. Moreover, the recognition of serotonergic neurons has been validated by an independent identification of the recorded neuron by its serotonin neuron specific expression of a fluorescent protein.

### 5.1. Comparison with existing procedures for serotonergic neuron identification from their physiological activity

Automatic routines for online measurements of action potentials can be designed, however until now no valid criteria for discriminating between spikes generated by serotonergic and non-serotonergic neurons have been established. Recognition of serotonergic neurons during extracellular recordings relies mostly on visual evaluation of the shape of the spike, that is often polyphasic, combined with the regular firing activity at relatively low frequency (up to 3-4 Hz). Thus, the mean criterion is mainly based on the asymmetric proportion between the upstroke and the downstroke of the spike (with a ratio usually *>*2.5) and duration of the spike itself after the main upstroke (usually accepted in the range *>* 1.2 ms). Coexistance of these characteristics is sufficient to enable an experienced Researcher to identify typical serotonergic neurons with a high degree of confidence. Nevertheless, in our recordings from genetically identified serotonergic neurons we noticed relatively frequent deviations from these criteria. Indeed, the spike duration of several neurons was less than 0.9 ms, down to 0.4-0.5 ms (see Fig. 6 in Mlinar et al., 2016). Similarly, a not negligible percentage of non-serotonergic neurons displayed spike shape and firing characteristics different from the expected biphasic, symmetric spikes of brief duration (*<*0.5-0.6 ms) and high frequency, often irregular, firing. Thus, some non-serotonergic neurons show long and asymmetric action potentials and sometimes have a regular, low frequency activity (*<*4 Hz) which makes their recognition difficult. Therefore, while “typical” serotonergic and non-serotonergic neurons are relatively easy to be discriminated with the currently accepted criteria a number of serotonergic neurons that do not comply with the classically established recognition criteria are discarded and not studied for their pharmacological and physiological characteristics.

Given these limits of the online visual recognition of serotonergic neurons, our model provides a valid tool for the intra-experiment identification of neurons recorded in the dorsal raphe nucleus, as the model can be implemented in the initial routine of *in vitro* recordings. Notably, our DL biological model relies only on spike shape for recognition of serotonin neurons and therefore it enables the identification and investigation of subpopulations of serotonergic neurons displaying irregular firing o low frequency oscillatory patterns of firing (see e.g. Mlinar et al., 2016).

### 5.2. Characteristics and limits of the model for its application

It is noteworthy that in several experiments ( *∼* 30% of those used here) used for training the model we applied a gentle suction in the patch pipette during the recording to improve the signal to noise ratio. We have previously shown (Mlinar et al., 2016) that this procedure does not alter the shape and duration of the recorded signals. In our context, this intra-experiment change in the amplitude of events recorded from the same neuron increases the robustness of the training data because implemented the model with events of constant shape but different weight of the background noise on the recorded signal. On the other hand, this was probably one source of the overfitting found in the initial, preliminary, model where the processed spike traces were longer (7 ms) than those used in the final models (4 ms). Indeed, in the presence of small and larger spikes with the same noise the DL processing could have retained the background noise as a signature of serotonergic neurons in addition to their shape and therefore this may explain the improvement of the model obtained by shortening the traces to be processed and limiting the recognition process to the spike shape. Notably, our synthetic model in which various background masks were superimposed to 4 ms spikes did not significantly improve the metrics compared to those of the biological model obtained using original 4 ms spikes, confirming that limiting the DL process to the spike was sufficient to eliminate the overfitting caused by the background recognition together with the spike shape for categorizing of neuron type.

It shoud be mentioned that the present model applies to the specific recording method used in collecting our database of spikes. Thus, for immediate application of the model the sampling frequency should be set at 40 kHz. Our data were acquired using the Clampex program in loose-seal cell attached patch clamp mode, but since the routine transforms the signals in *.csv files any acquisition program that produces files in a format that can be transformed in *.csv format would provide adequate input for the model. The amplitude of the recorded current should be greater than the detection threshold that we have imposed in the model to minimize acquisition of small transients (*>*50 pA). Finally, our recordings were performed at the temperature of ~37 °C. Although small deviations from this temperature could be tolerated, it should be considered that the width of the spike, may be influenced by temperature. Notwithstanding these limitations, if the sampling rate is adequate and the signal reaches the detection amplitude the model provides an answer on neuron type with an accuracy of *>*88.4% within an inference time of a few milliseconds after the submission of the recorded traces (the inference time is the raw time taken by the model in classifying the signal without considering the latent time of converting and transmitting the signal to the model which can vary depending on the user interface chosen in the deployment of the model).

*Perspectives.* Importantly, a relatively low number of recordings was sufficient to develop our deep-learning based model. In perspective, the procedure we describe can be applied to construct further models for the identification of other spontaneously active monoaminergic neurons. For instance, our approach with genetically fluorescent mice can be extended to the recognition of other neurons for *in vitro* recordings. Similarly, application of the CNN deep-learning procedure to neuronal types recognized with optogenetic methods (Liu et al., 2014) or with post-hoc immunohistochemistry (Allers and Sharp, 2003) *in vivo* may enable to construct a template of models capable to recognize a variety of neurons during *in vivo* recordings from mouse and rats. Once validated, these models would allow rapid identification of the recorded neuron, making *in vivo* recording of the activity of selected neurons more feasible and less demanding than at present. This may also facilitate studies on the correlation between the firing of different neuron types and behavioural responses in laboratory animals and increase our understanding of the physiological role of these neurons in modulating higher brain functions. In conclusion, our model provides the first proof of concept that neurons can be recognized from the sole characteristics of extracellularly recorded spikes and independently of their firing rhythm. Our model could readily be applied for intra-experiment decision making on the experimental design to apply to record that specific neuron and/or for helping the training of young Researchers at the beginning of their experience.

## 6. Acknowledgments

The original recordings and measurements of the spikes were performed by Dr. Boris Mlinar and Dr. Alberto Montalbano.

## 8. Data Sharing Statement

The data and source code correspondent to the analyses contained in this manuscript are publicly available from: Corradetti et al. 2024 Repository with all data available at github.com/neuraldl/DLAtypicalSerotoninergicCells.git

## 9. Author Contributions

All authors have made a significant contribution to the idea formation, study design, data curation, analysis and interpretation. D.C. & R.C. wrote and reviewed the manuscript. R.C. collected and selected the data. A.B. and D.C. realised the software for both the neural networks and the synthetic data generation.

## Notes

https://github.com/neuraldl/DLAtypicalSerotoninergicCells

### Competing Interest Statement

The authors have declared no competing interest.

